# Triple dissociation of attention and decision computations across prefrontal cortex

**DOI:** 10.1101/171173

**Authors:** LT Hunt, WMN Malalasekera, AO de Berker, B Miranda, SF Farmer, TEJ Behrens, SW Kennerley

## Abstract

Anatomical^1,2^, neuroimaging^3,4^ and lesion studies^5,6^ indicate that prefrontal cortex (PFC) can be subdivided into different subregions supporting distinct aspects of decision making. However, explanations of neuronal computations within these subregions varies widely across studies^7-22^. An integrated and mechanistic account of PFC function therefore remains elusive. Resolving these debates demands a rich dataset that directly contrasts neuronal activity across multiple PFC subregions within a single paradigm, whilst experimentally controlling factors such as the order, duration and frequency in which choice options are attended and compared. Here, we contrast neuronal population responses between macaque orbitofrontal (OFC), anterior cingulate (ACC) and dorsolateral prefrontal cortices (DLPFC) during sequential value-guided information search and choice. From the first fixation of choice-related stimuli, a strong triple dissociation of information encoding emerges in parallel across these PFC subregions. As further information is gathered, population responses in OFC reflect an attention-guided value comparison process. Meanwhile, parallel signals in ACC reflect belief updating in light of new evidence, integration of that evidence to a decision bound, and an emerging action plan for which option should be chosen. Our findings demonstrate the co-existence of multiple, distributed decision-related computations across PFC subregions during value-guided choice. They provide a synthesis of several competing accounts of PFC function.

---

Recent debates on the role of PFC subregions in value-guided decision making have included whether PFC acts in serial (certain subregions preceding others) or parallel (simultaneous, distributed activity across subregions)^7,8^; whether and how attention shapes the decision process, in particular in OFC^9-13^; and whether ACC integrates evidence for different actions^14-17^, modifies behaviour in light of new evidence^18-20^, or evaluates evidence for alternative courses of action^21,22^.

We addressed all of these questions by recording activity from ACC, DLPFC and OFC (n=189, 134 and 183 single neurons respectively, see **Extended Data Figure 1** and **Methods** for details) during a decision task performed by two macaque monkeys (*M. mulatta*). Our task design **(Figure 1a)** mirrored behavioural studies examining attention-guided information search during sequential, multi-attribute choice^23,24^. Each *option*, presented on left and right sides of the screen, comprised two pre-learned picture cues, representing different *attributes*, probability and magnitude of juice reward. Crucially, at trial start, all cues were hidden. Subjects made an instructed saccade towards a highlighted location to reveal cue 1. Following 300ms uninterrupted fixation, cue 1 was recovered and another location highlighted, either vertically on the same *option*, or horizontally on the same *attribute.* Subjects saccaded here to reveal cue 2, again for 300ms. Hereafter, subjects could choose either option using a manual left/right joystick movement. Alternatively, they could fixate one or both remaining highlighted cues in any order to reveal further information before committing to their choice. Following joystick choice, all four cues were revealed and juice reward was delivered with the chosen probability and magnitude. Picture cues, first/second highlighted location, and probability/magnitude attribute on top vs. bottom were pseudorandomly selected on each trial (with uniform distribution).

**Figure 1.**
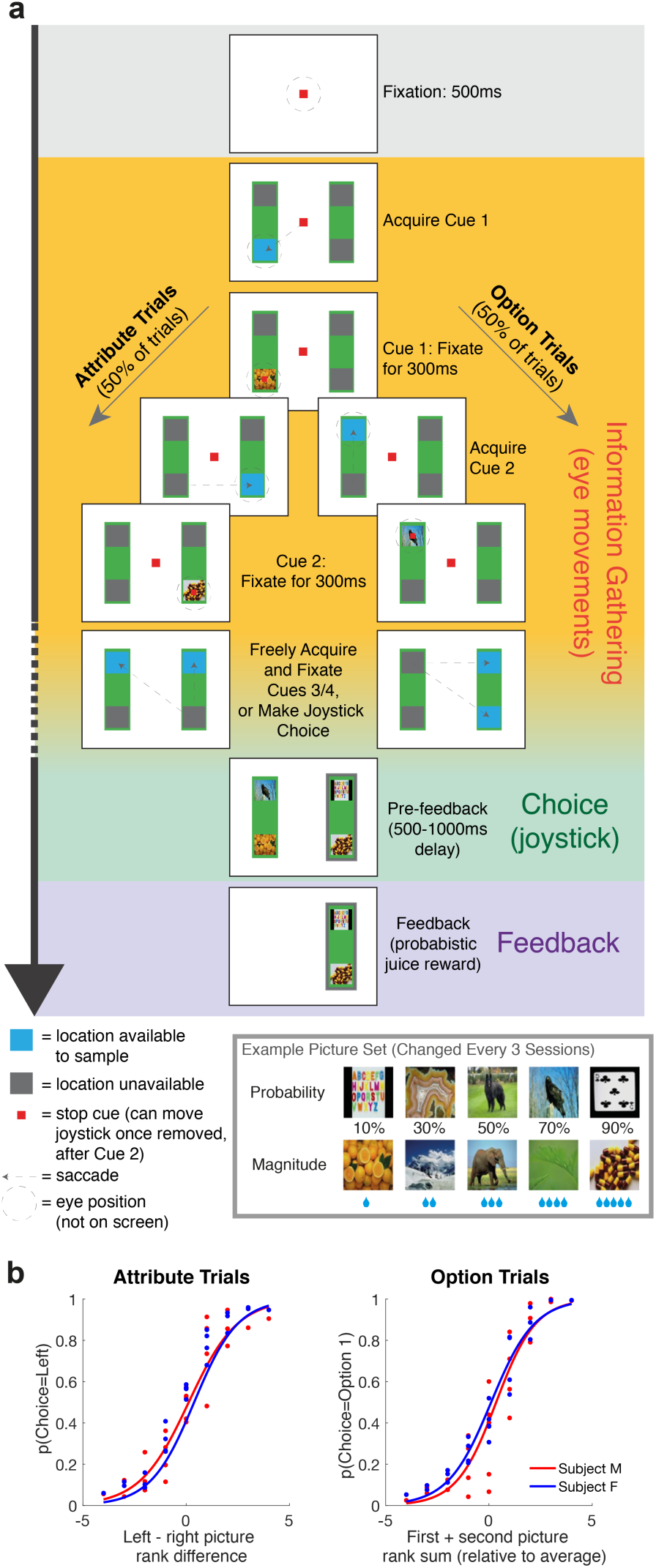
Experimental paradigm and basic subject behaviour. **(a)** Task design. Subjects chose between a left and right option (green rectangles) using a manual joystick movement, after sequentially sampling 2, 3, or 4 cues that revealed reward probability and magnitude to the subject. Blue squares indicated locations available for information sampling. On attribute trials (left panels), cues 1 and 2 were on opposing options but the same attribute; on option trials (right panels), cues 1 and 2 were different attributes of the same option. **(b)** Choice behaviour as function of cue 1 and 2 value. **(i)** Probability of choosing left option on attribute trials, as function of left-right picture rank difference (where 1 is lowest rank picture on each attribute, and 5 is highest rank picture). **(ii)** Probability of choosing option 1 on option trials, as function of first plus second picture value (relative to middle value picture 3). See also **Extended Data Figure 2** and **Supplementary Note 1** for additional analyses of choices/information sampling.

Both monkeys used cue values appropriately to guide their choices **(Figure 1b)**. They chose the option with higher expected value on 77.0% and 80.1% of trials (monkeys F (n=25 sessions) and M (n=32 sessions) respectively), assigning approximately equal weight to both reward probability and magnitude, and using all viewed cues to guide their choice **(Extended Data Figure 2)**. However, most choices were based upon partial information: subjects chose before all four cues had been evaluated on 85.5%/71.4% of trials (subjects F/M respectively) (see **Supplementary Note 1** for further discussion of information sampling behaviour).

A critical feature of our task design is that the cue that is currently being attended can be decomposed into multiple features: its associated *action* (left/right joystick response required to choose that option), *attribute* (magnitude or probability), *spatial position* (presented on top/bottom of screen), and *value* (level of reward probability/magnitude). In line with previous studies, we found a degree of PFC subregion specificity in single neuron encoding of these features. However, there was also substantial between-neuron *heterogeneity* of decision variable encoding within each subregion **(Supplementary Note 2)**.

We capitalised upon this response heterogeneity by assessing population-level encoding of decision variables. At cue 1, we used representational similarity analysis (RSA). RSA correlates the normalised firing rate of the neural population between all conditions of interest^25^. This characterises task encoding across the neural population without strong prior assumptions on its structure. Here, we consider 20 such conditions: 5 probability cues and 5 magnitude cues, presented on either the left or right option.

Using this approach we examined whether decision-related information emerged sequentially or in parallel at cue 1 presentation^7,8^. RSA revealed distinct task-evoked neural codes within each subregion **(Figure 2a-c)** that were consistent across subjects **(Extended Data Figure 3)** and emerged largely in parallel **(Extended Data Movie 1)**. To formally compare subregion specificity and temporal evolution of population representations, we regressed templates onto RSA matrices to capture different features of the task design. DLPFC RSA reflected whether the subject was attending left or right **(Figure 2d)**; OFC reflected the currently attended stimulus identity, irrespective of spatial postition **(Figure 2e)**; OFC representational similarity was also high for cues of similar attended value **(Figure 2f)**; ACC and DLPFC RSA showed a value code modulated by left/right action **(Figure 2g)**. For intuition, we provide single neuron examples for each of these features in **Extended Data Figure 4**. We also present RSA matrices subdivided by top/bottom spatial position in **Extended Data Figure 5**.

**Figure 2.**
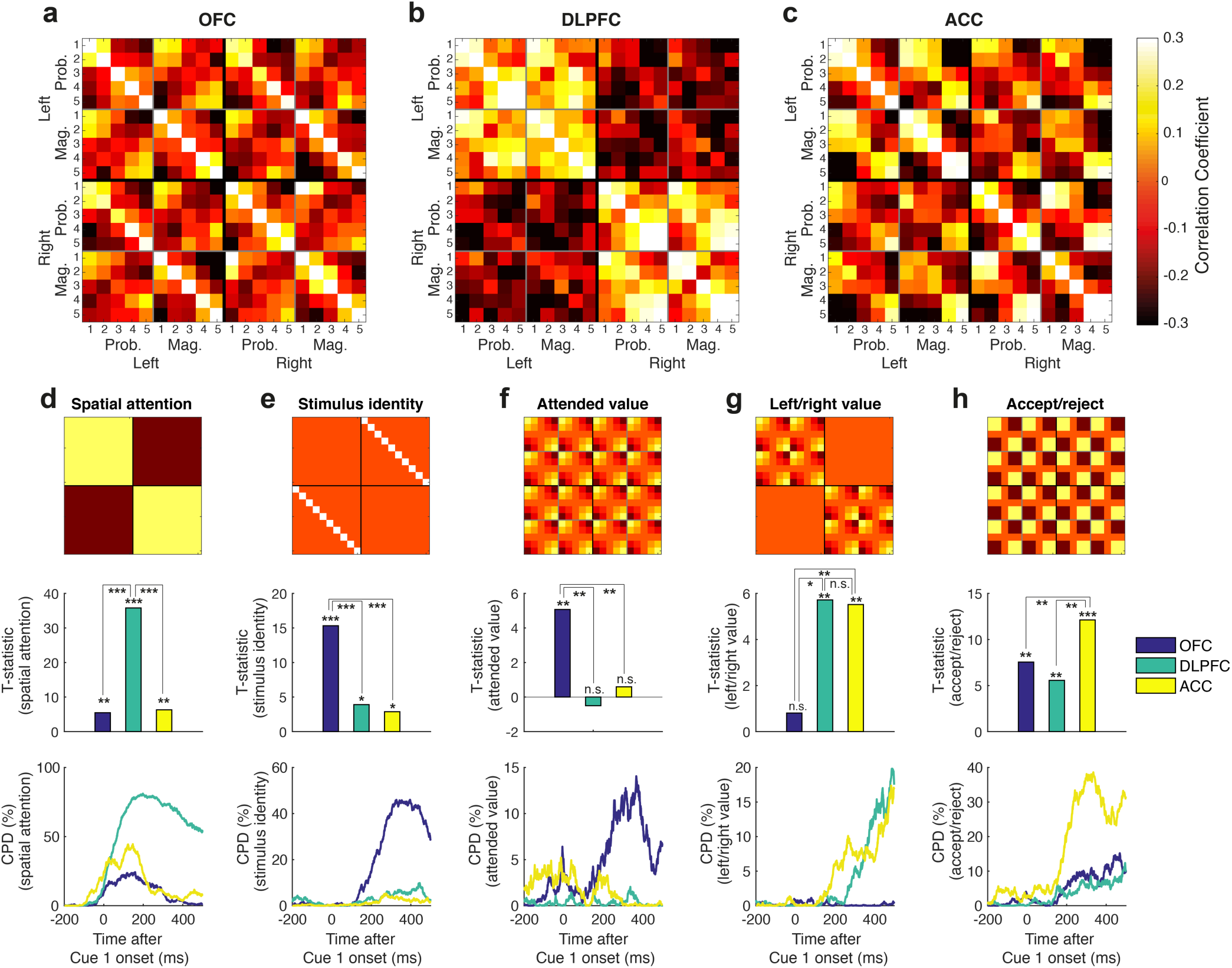
Triple dissociation of task-evoked neural codes across OFC, DLPFC and ACC at Cue 1 presentation. **(a)** OFC representational similarity analysis between the 10 different cue identities – probability and magnitude, sorted from lowest (1) to highest (5) ranked picture – when presented on left and right options. For matrix element (*i*,*j*), colour denotes correlation coefficient of Z-scored firing rate between condition *i* and condition *j* across the OFC neuronal population. Firing rate is averaged from 100ms to 500ms poststimulus (see also **Extended Data Movie 1)**. Note this matrix collapses across top/bottom spatial position; see **Extended Data Figure 5** for 40x40 matrix, splitting conditions by spatial position. **(b)/(c)** As panel **(a)**, but for DLPFC and ACC populations respectively. **(d)-(h)** Results from multiple linear regression of ‘templates’ onto RSA matrices. In each column, top subpanel shows template matrix, middle subpanel shows corresponding regression T-statistics for each region, and bottom subpanel shows coefficient of partial determination from sliding regression of templates onto RSA matrices, using sliding window of +/- 100ms. See **Methods** for full description of template matrices, regression model and statistical inference via non-parametric permutation test. **(d)** ‘Spatial attention’ template (differentiating cues on left vs. right-hand option) was particularly prominent in DLPFC (DLPFC: T_399_=35.782, p<1^∗^10^−4^), and significantly more so than other regions (one-way ANOVA F_2,1179_ = 311.18, p<1^∗^10^−4^; post-hoc comparison for DLPFC>ACC: p<1^∗^10^−4^; post-hoc comparison for DLPFC>OFC: p<1^∗^10^−4^); **(e)** ‘stimulus identity’ template (responding similarly to the same cue irrespective of side) was particularly strong in OFC (T_399_=15.3173, p<1^∗^10^−4^), again more so than other regions (F_2,1179_ = 32.77, p<1^∗^10^−4^; post-hoc comparisons: OFC>ACC: p<1^∗^10^−4^; OFC>ACC: p<1^∗^10^−4^); **(f)** ‘Attended value’ template (representing cue 1 value irrespective of stimulus location or attribute) was prominent in OFC (T_399_ = 5.0697, p=0.0036), and significantly more so than in other regions (F_2,1179_ = 6.73, p=0.0126; post-hoc comparisons: OFC>DLPFC: p=0.0017; OFC>ACC: p=0.0002); **(g)** ‘Left/right value’ template was prominent principally within ACC (T_399_ = 5.5 1 51, p=0.0096) and DLPFC (T_399_=5.7156, p=0.0062), and significantly more so than in OFC (F_2,1179_ = 9.08, p=0.011; post-hoc comparisons: ACC>OFC: p=0.0003; DLPFC>OFC: p=0.011); **(h)** ‘Accept/reject’ template (reflecting whether Cue 1 had high (rank 4 or 5) versus low value (rank 1 or 2)) was strongest in ACC (T_399_=12.1217, p<1^∗^10^−4^) and significantly more so than other regions (F_2,1179_ = 17.20, p=0.0006; post-hoc comparisons: ACC>OFC: p<1^∗^10^−4^; ACC>DLPFC: p=0.0004).

Notably, RSA *divided* high-valued and low-valued items, particularly in ACC **(Figure 2c)**. Variance in ACC was therefore best explained as a non-linear, step-like function of value **(Figure 2h)**. This contrasted with the OFC’s linear representation of cue 1 attended value **(Figure 2a/f)**. The ACC signal may have a similar interpretation to ACC activity signalling shifts in belief or behaviour across multiple trials^18,20-22^ – with an ‘accept’ signal being akin to *confirming* the currently attended option, but the ‘reject’ signal being akin to *disconfirming* it. The OFC signal, by contrast, might be a natural substrate to support comparison of *currently attended* cue value versus previously attended (*stored*) cue values^10,12^.

We tested these ideas further by investigating how population encoding of value evolved across time as further cues were uncovered. We used multiple linear regression to evaluate how strongly each neuron encoded the values of cues 1, 2, 3 and 4 across time, on both option and attribute trials. In **Figure 3a**, we plot the average coefficient of partial determination (CPD, a measure of variance explained by each regressor; see **methods)**, timelocked to each of the first three cues. This shows that value encoding by OFC neurons peaked approximately 300ms after each cue was presented, but was then sustained above baseline as further cues were attended.

**Figure 3.**
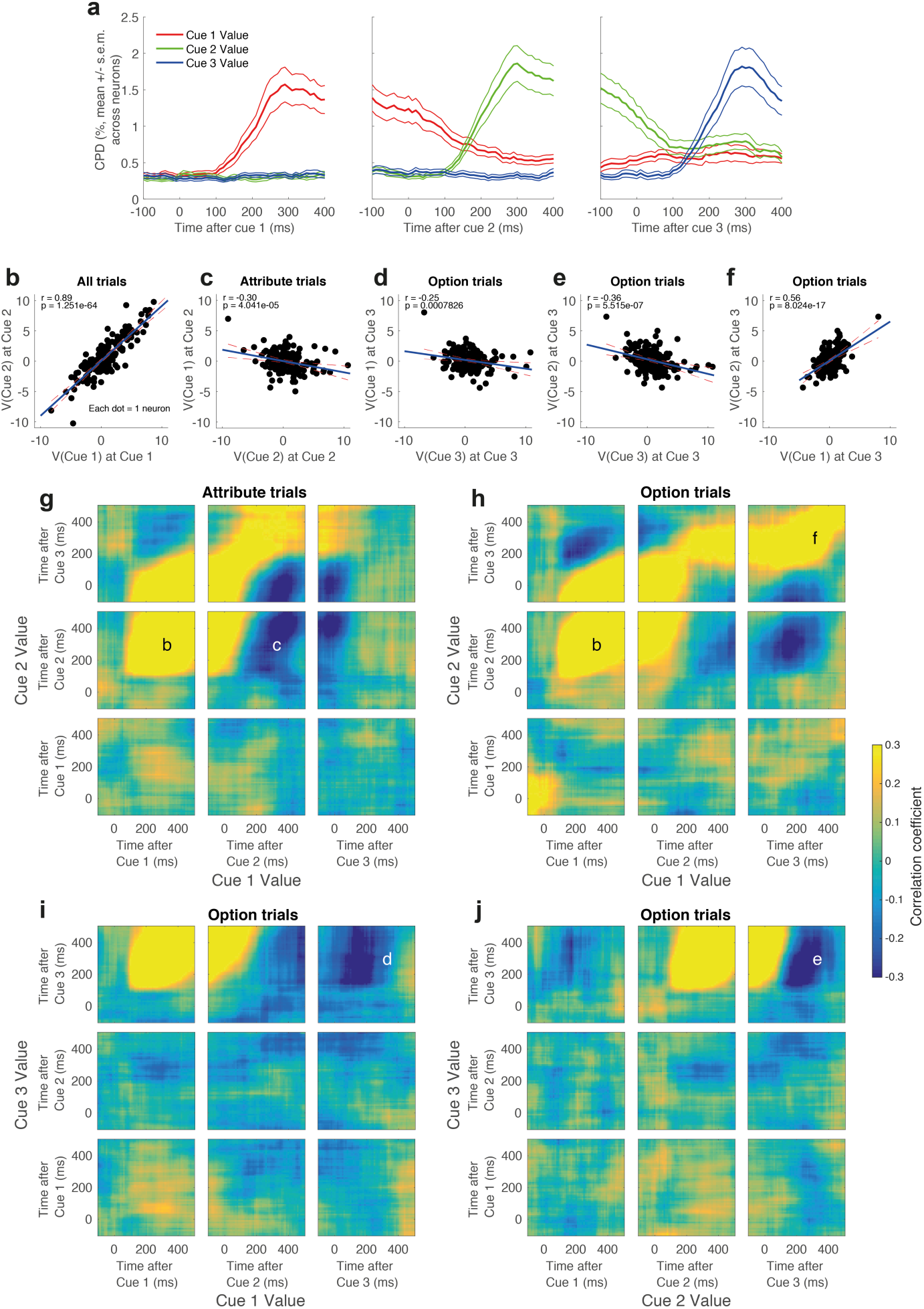
Valuation subspaces for attended (‘online’) and stored cues supporting attention-guided value comparison in OFC. **(a)** Coefficient of partial determination for Cue 1 Value, Cue 2 Value and Cue 3 value, timelocked to each cue’s presentation using a sliding 200ms window. Lines denote mean +/- s.e.m. across neurons. **(b)** Positive relationship between regression coefficients for value of cue 1 when cue 1 is presented (ordinate), and value of cue 2 when cue 2 is presented (abscissa), implying a stable subspace reflecting the currently attended value. **(c)** When cue 2 is being attended on attribute trials, the online value subspace (ordinate) correlates negatively with the subspace reflecting the stored value of cue 1 (abscissa). **(d)-(f)** When cue 3 is being attended on option trials, the attended value subspace (ordinate of **(d)** and **(e)**) correlates negatively with the subspace for stored values of both cue 1 (abscissa of **(d)**) and cue 2 (abscissa of **(e)**). The two stored subspaces are positively correlated **(f)**. **(g)-(i)** Cross-correlation matrices reflecting the time-varying relationship between different value subspaces on attribute trials (in **(g)**) and option trials (in **(h)-(i)**). Heatmaps reflect the correlation coefficient between regression coefficients for each cue’s across the OFC population. Superimposed letters denote the correlations that are plotted above, in panels **(b)-(f)**. See also **Extended Data Figures 6-7**.

Crucially, neuronal regression coefficients were highly variable across the population. We again capitalised upon this heterogeneity to define population ‘subspaces’ for value encoding^12,26^. For example, regression coefficients for cue 1 value when cue 1 was attended (ordinate in **Figure 3b)** correlated positively across neurons with regression coefficients for cue 2 value when cue 2 was attended (abscissa in **Figure 3b)**. These two regressors are orthogonal and defined at different task epochs. This analysis therefore reveals a stable population subspace for the *currently attended* cue value.

We repeated this approach for different phases of the task, to ask how the *currently attended* cue value subspace (ordinates in **Figures 3c-e)** correlated with subspaces encoding previously attended, or *stored*, cues (abscissae in **Figures 3c-e)**. For example, when cue 2 was attended on attribute trials, the currently attended cue 2 value subspace correlated *negatively* with the stored cue 1 value subspace, representing the other option **(Figure 3c)**. When cue 3 was attended on option trials, the *stored* cue 1 and *stored* cue 2 values both represented the other option, and were both negatively correlated with *currently attended* cue 3 value subspace **(Figure 3d/e)**. However, these two stored subspaces were themselves positively correlated on option trials **(Figure 3f)**. Collectively these results reveal the signature of attention-guided value comparison in OFC between the currently attended cue and previously attended cues stored in working memory.

A more complete description of the interaction between attention and value across time can be obtained by plotting the cross-correlation of these subspaces **(Figure 3g-j)**. The letters superimposed on these plots refer back to the correlations shown in **Figure 3b-f**. This approach also reveals other key features of the relationship between value subspaces in OFC, at other timepoints. Notably, the signature of attention-guided value comparison observed in these analyses was unique to OFC. Whilst the *currently attended* value subspace was present in DLPFC/ACC, value comparison with *stored* cues was substantially weaker or absent in these regions **(Extended Data Figures 6-7)**.

We then evaluated ACC population activity across cues 2 and 3, guided both by a recent literature on ACC adapting behaviour in light of new evidence^18-20^, and our interpretation of **Figure 2h** as a potential *belief confirmation* signal. To test this hypothesis more rigorously, we included four regressors in our regression model that capture *belief confirmation* at subsequent cues, on both option and attribute trials. In particular, whenever the evidence presented to the subject thus far suggests that the *currently attended* side should be chosen, then ‘belief confirmation’ scales *positively* with value. Whenever the evidence suggests that the *unattended* side should be chosen, ‘belief confirmation’ scales *negatively* with value (see **Extended Data Figure 8)**. This means that all four regressors were thus orthogonal to *currently attended* value.

We used these regressors to test whether ACC population subspaces reliably encoded *belief confirmation.* We found that ACC population subspaces for each of these regressors were significantly correlated with each other, and also to cue 1 *belief confirmation* **(Figure 4a**/**Extended Data Figure 9)**. As all five regressors are orthogonal to each other and defined at different parts of the trial, this reveals a stable population code in ACC for belief updating as a decision unfolds.

**Figure 4.**
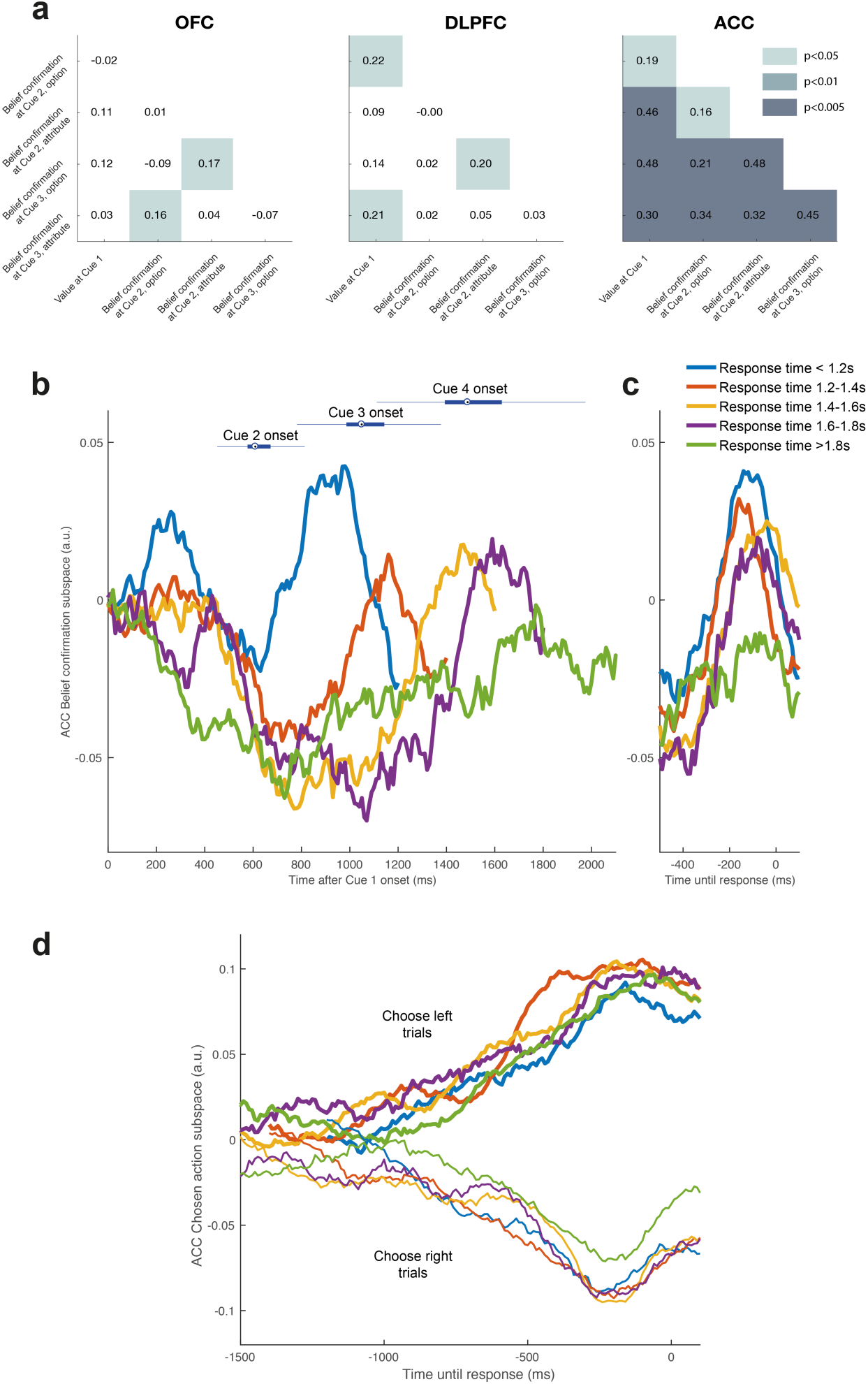
Multiple signals in ACC reflect belief confirmation, commitment to a course of action, and accumulation of evidence for left/right movement. **(a)** Positive population correlations between four orthogonal regressors that reflect ‘belief confirmation’ at cue 2/cue 3 on both option and attribute trials (cf. **Extended Data Figure 8**), and initial value population response at cue 1 (cf. **Figure 2h**), demonstrating a stable belief confirmation subspace in ACC across multiple cues. Shading denotes significant correlation of parameter estimates across the neural population in each region. See **Extended Data Figure 9** for individual ACC correlations. **(b)/(c)** Projecting ACC population activity into this subspace reveals ramping immediately prior to the commitment to joystick movement. In part **(b)**, trials are sorted by response time and time-locked to cue 1 onset (blue bars above are box plots of cue onset times for other cues); in part **(c)**, trials are time-locked to response onset. **(d)** An orthogonal subspace in ACC reflects the emergence of an action plan to choose the left or right option. Activity in this subspace progressively favours one action over the other across time. See **Extended Data Movie 2** for relationship between belief confirmation and action selection subspaces.

We next asked whether this belief confirmation subspace might support commitment to a final decision^15,17^. To answer this, we examined the temporal evolution of belief confirmation subspace activity, using the regressors in **Extended Data Figure 9**/**Figure 4a**. We used one half of the data to define the subspace, and projected the data from the remaining half into this subspace to examine its evolution across time. Time-varying activity within this subspace showed distinct dynamics on trials of different reaction times **(Figure 4b)**. However, it ramped towards a common threshold shortly prior to joystick movement, except on long reaction time trials **(Figure 4c)**. Activity within this subspace thus becomes prominent immediately prior to commitment to action.

Finally, **Figure 2g** and previous studies indicate that ACC contains a signal related to *action selection*^15-17,27^. We therefore defined a separate subspace for whether the animal would choose left or right on the current trial, adopting the same split-half approach as in **Figure 4b/c**. Activity in the ACC action selection subspace also gradually ramped as evidence was revealed about which option to choose, and peaked immediately prior to action selection **(Figure 4d)**. Note that ACC belief confirmation and action selection subspaces are orthogonal to one another; the relationship between them can be seen in **Extended Data Movie 2**. This finding therefore supports the recently expressed notion of multiple complementary signals in ACC during choice^28^.

Despite evidence that PFC subregions make dissociable contributions to value-based choice^3-6^, uncovering how PFC dynamically compares current versus past evidence and determines when to commit to a choice has remained unclear. Several key computations must occur, including stimulus identification, valuation and integration with other attributes, comparison to previous stimuli and action selection. It has remained contentious whether these computations occur in serial or in parallel and how they are distributed across brain areas^7,8^. Our findings indicate parallel emergence of subregion-specific computations from the very first fixation of decision information **(Figure 2)** and distinct computations within OFC and ACC leading to choice **(Figures 3/4)**.

In real-world decision tasks, value-guided decision-making is shaped heavily by attention. Information gathering strategies of both human consumers^9,23,29^ and foraging animals^30^ are well characterised as consecutive consideration of each alternative and its component attributes. We found that DLPFC represented where subjects’ spatial attention was currently being deployed **(Figure 2d)** whereas OFC reflected an attention-guided value comparison **(Figure 3)**. Yet neurophysiological studies of decision-making often neglect information search, and prevent free viewing of choice information by requiring central fixation (with notable exceptions^10,12^). Being unable to monitor covert attention as it varies across trials may confound the isolation of the individual computations of the decision process, as revealed in the present study. Further integrating attention-guided information search and choice may prove key in extending findings from neuroeconomics to more naturalistic decision scenarios, where options are evaluated and compared in a consecutive fashion.

---

**Extended Data Figure 1.**
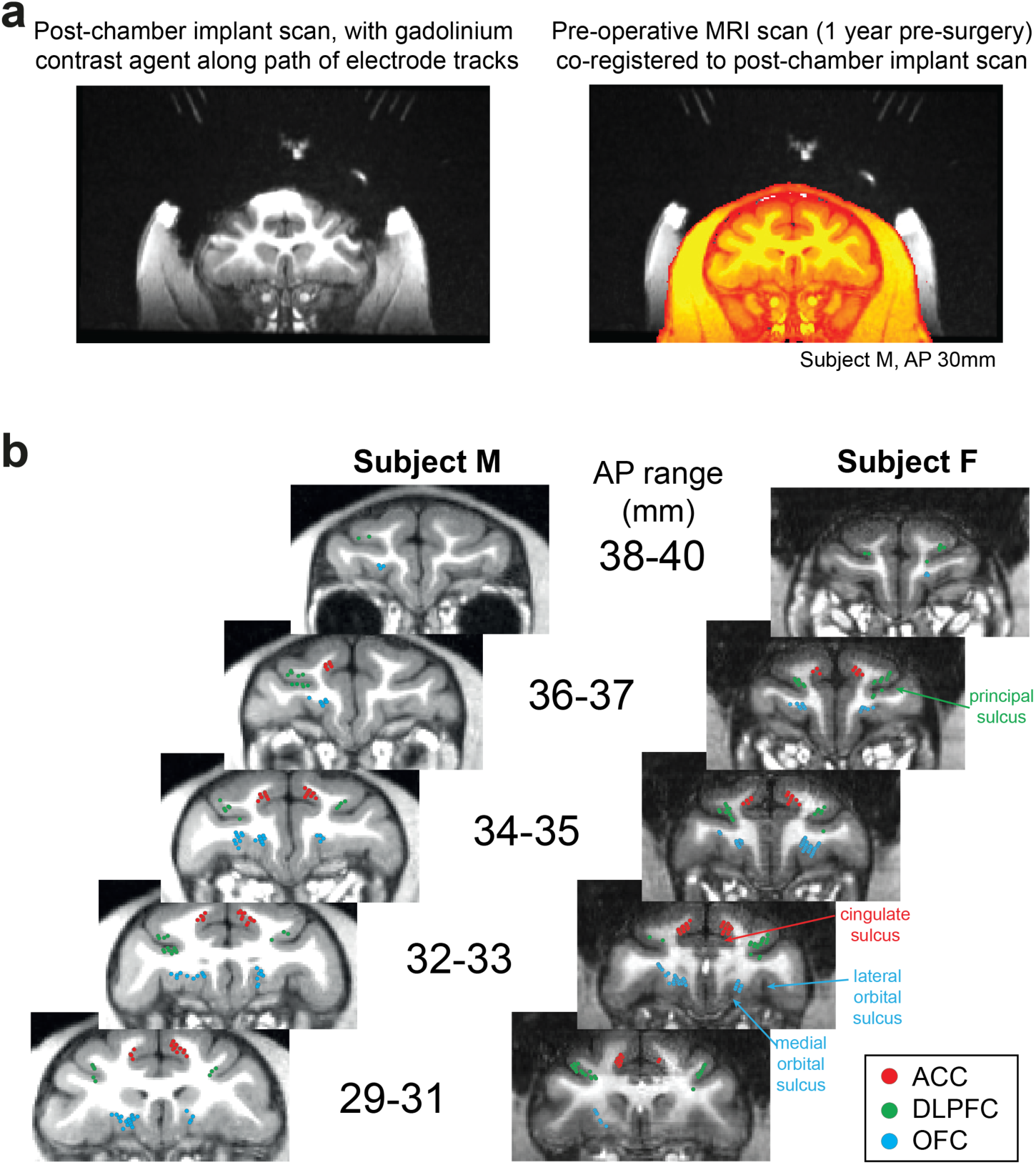
Recording locations. (a) Strategy for reconstructing path of electrode tracks. Left: After surgery for chamber implantation, prior to craniotomy, subjects underwent MRI scan with a custom-built implant placed within the chamber. This contained the MRI contrast agent gadolinium along the trajectory of potential recording paths at regular 4mm intervals. Note the prominent susceptibility artefact due to titanium chambers does not affect gadolinium trajectories, which were intentionally located away from the chamber. Right: This scan could be co-registered to a preoperative scan without susceptibility artefact (in orange; note that head appears smaller due to muscle growth between scans), to reliably reconstruct recording locations. This technique was further verified by identification of grey/white matter boundaries during lowering of electrodes along different trajectories. **(b)** Recording locations of orbitofrontal neurons (OFC), dorsolateral prefrontal neurons (DLPFC), and anterior cingulate neurons (ACC), shown on coronal sections. AP range denotes position anterior to interaural plane in stereotactic coordinates.

**Extended Data Figure 2.**
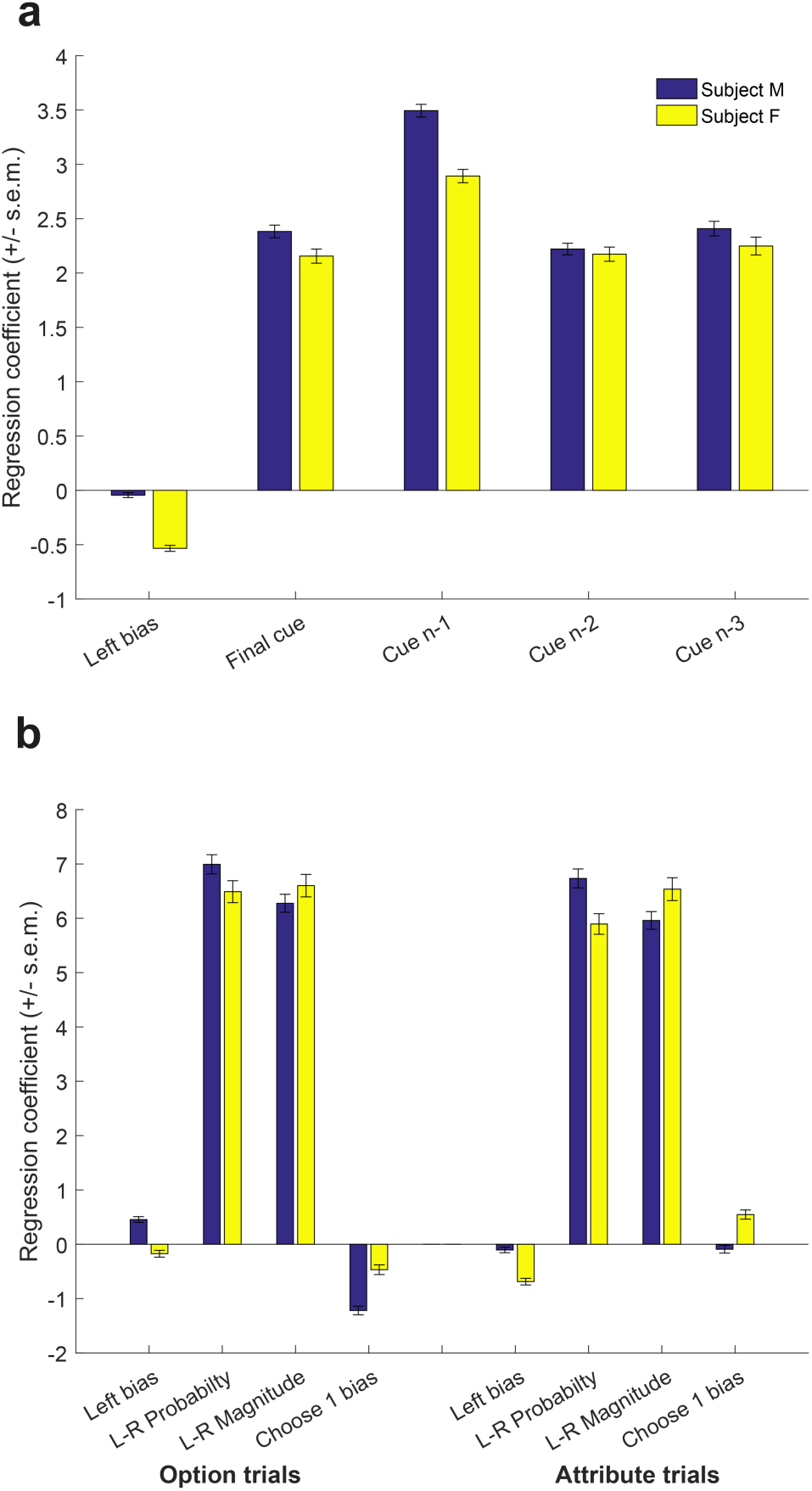
Logistic regression shows weighting of all four presented cues and both probability and magnitude attributes on subject choices. **(a)** Bars show mean +/- s.e.m. of regression coefficient for the final viewed picture, penultimate viewed picture (n-1), antepenultimate viewed picture (n-2), and first viewed picture on trials where subject viewed all four cues (n-3), as well as bias towards choosing the left cue. Both subjects use all four cues to guide their choices, with a slight upweighting of the antepenultimate cue. **(b)** Bars show mean +/- of the regression coefficient for left minus right probability, and left minus right magnitude, separately for option and attribute trials. Also included in the model are a bias towards choosing the left cue and a bias towards choosing the first side (note that both subjects show a small but significant bias towards choosing the first side on option trials, also visible in main **Figure 1b**).

**Extended Data Figure 3.**
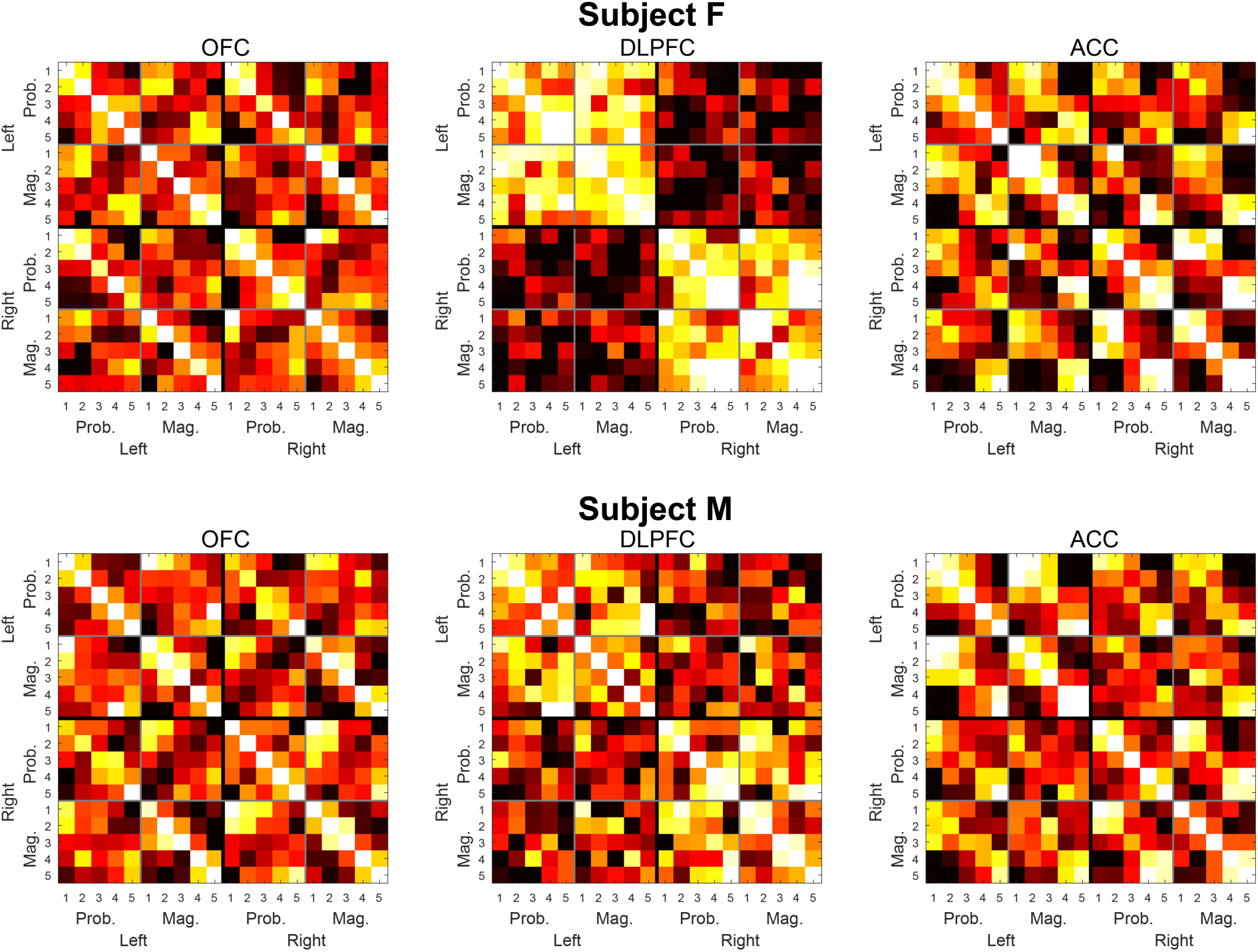
Reproducibility of representational similarity matrices across subjects. Data are as presented in main **Figure 2a-c**, plotted separately for subjects M and F.

**Extended Data Figure 4.**
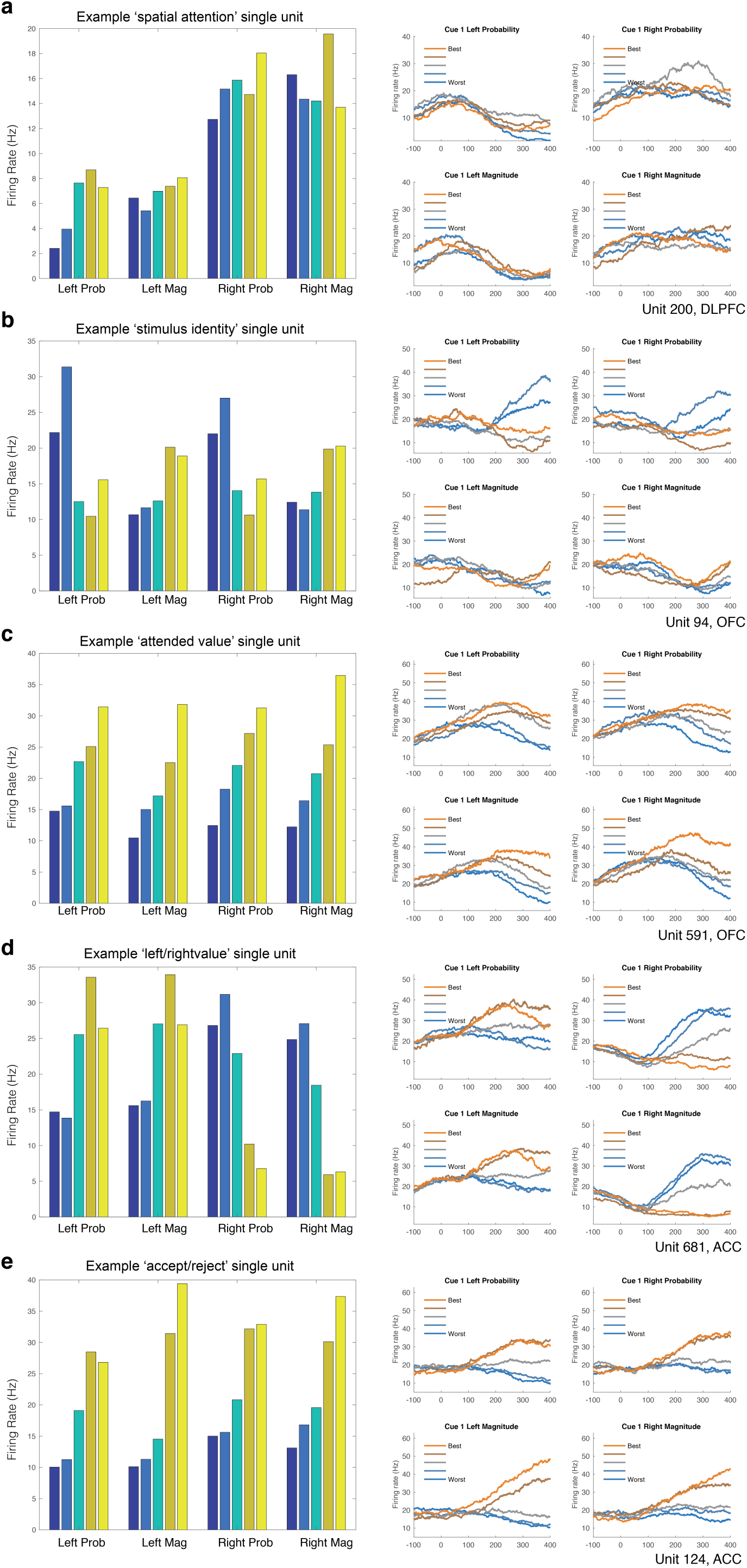
Example single neurons to provide intuition for how different task variables are represented at cue 1 presentation. Each subplot shows a different single neuron example. Bar plots show average firing rate of the neuron to each of the 10 stimuli when presented on left and right of the screen, averaged from 300-500ms following cue 1 presentation. Blue colours are lowest value (worst) stimuli, yellow colours are highest value (best) stimuli. Line plots show peri-stimulus histograms for each of these conditions, timelocked in ms to cue 1 presentation. **(a)** Example DLPFC unit reflecting spatial position (high firing for all stimuli on right side of screen. **(b)** Example OFC unit showing ‘stimulus identity’ coding (esoteric high firing for certain stimuli, replicated on both left/right sides of screen). **(c)** Example OFC unit showing ‘attented value’ coding (firing linearly scales with value, irrespective of attribute or spatial position). **(d)** Example ACC unit reflecting action value (high firing for high valued stimuli on left or low valued stimuli on right). **(e)** Example ACC unit showing ‘accept/reject’ coding (high firing for stimuli ranked 4 or 5; low firing for stimuli ranked 1 or 2).

**Extended Data Figure 5.**
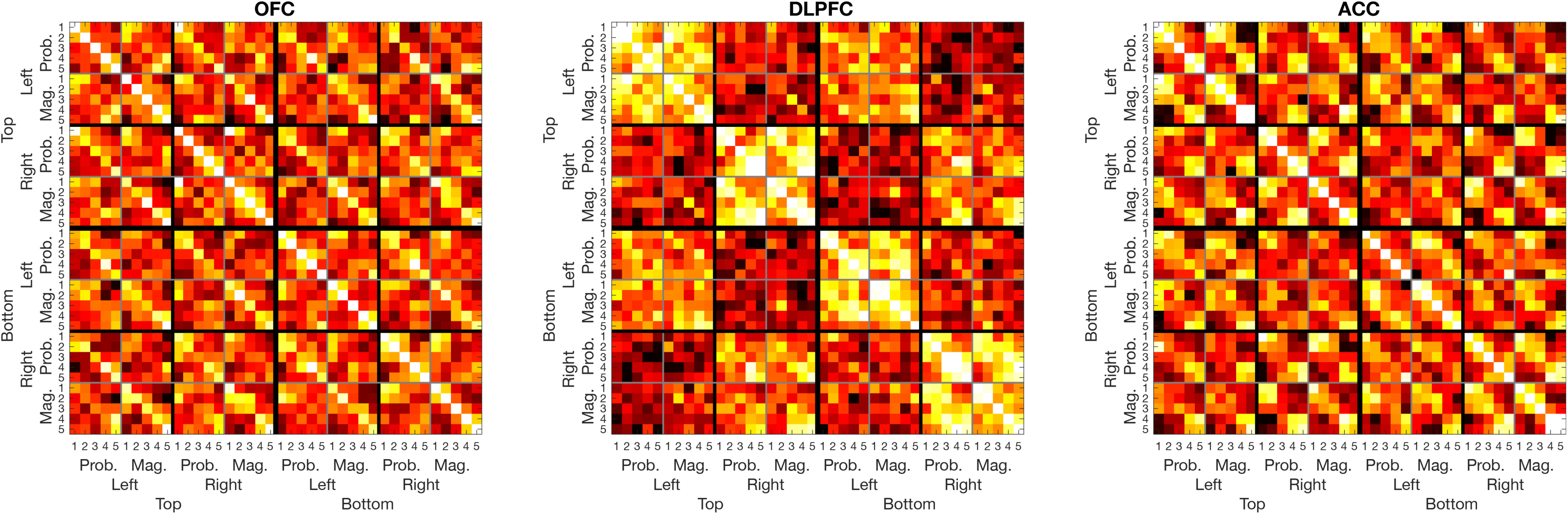
Representational similarity across all 40 conditions at cue 1 presentation. Data are as presented in main **Figure 2a-c**, but are now subdivided into trials where cue 1 was presented in the top versus bottom half of the screen. The key results from this analysis primarily replicate the findings when top and bottom cues are collapsed. Note, however, that in DLPFC, representational similarity is modulated by top/bottom stimulus location (compare, for example, the average brightness for top left→top left versus top left → bottom left). This indicates that DLPFC activity primarily represents spatial position rather than simply left/right action. This finding was also recapitulated in single unit analyses of DLPFC neurons (see **Supplementary Note 2**).

**Extended Data Figure 6.**
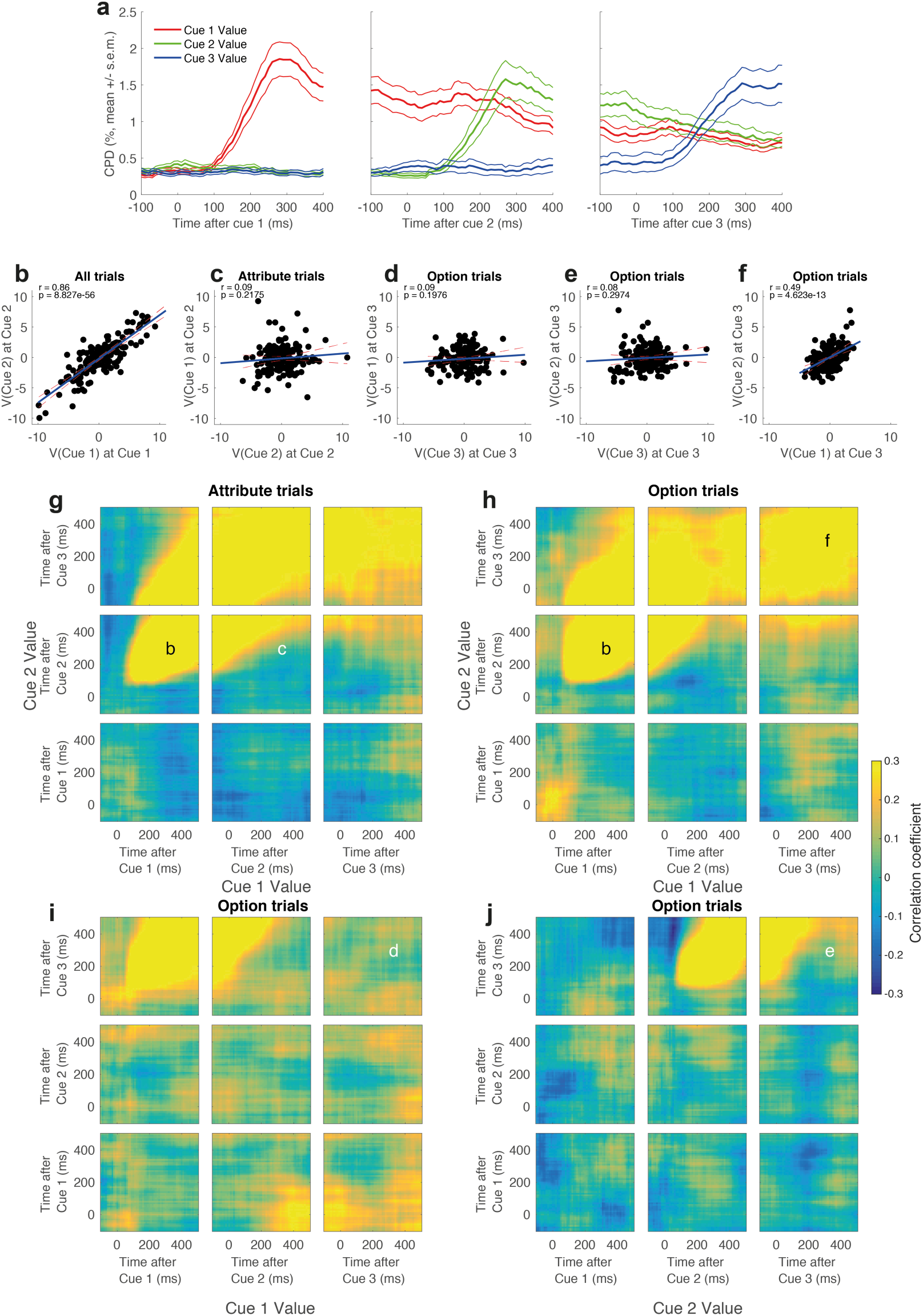
Value subspaces for attended and stored code in ACC. Figure layout is as for main **Figure 3**. Note that like OFC, ACC shows consistent single neuron coding of value **(a)**, and that population subspaces for attended **(b)** and stored **(f)** subspaces are present in ACC. However, there is no evidence of inhibition between the attended and stored subspaces (parts **(c)-(e)**).

**Extended Data Figure 7.**
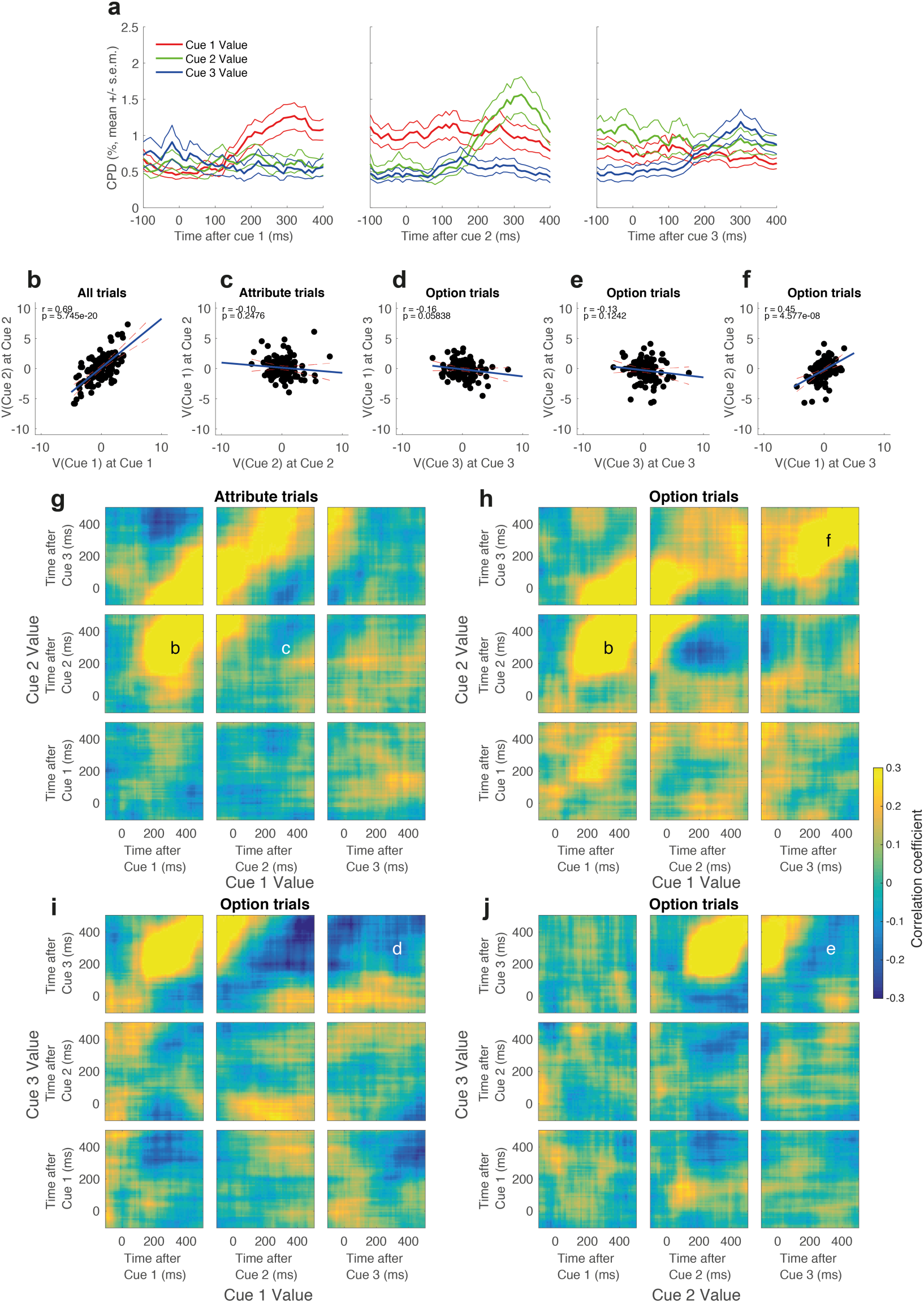
Value subspaces for attended and stored code in DLPFC. Figure layout is as for main **Figure 3**. Value coding is weaker in DLPFC than in either of the other two regions **(a)**, but population subspaces for attended **(b)** and stored **(f)** subspaces are nonetheless present in DLPFC. However, there is little evidence of inhibition between the attended and stored subspaces (parts **(c)-(e)**).

**Extended Data Figure 8.**
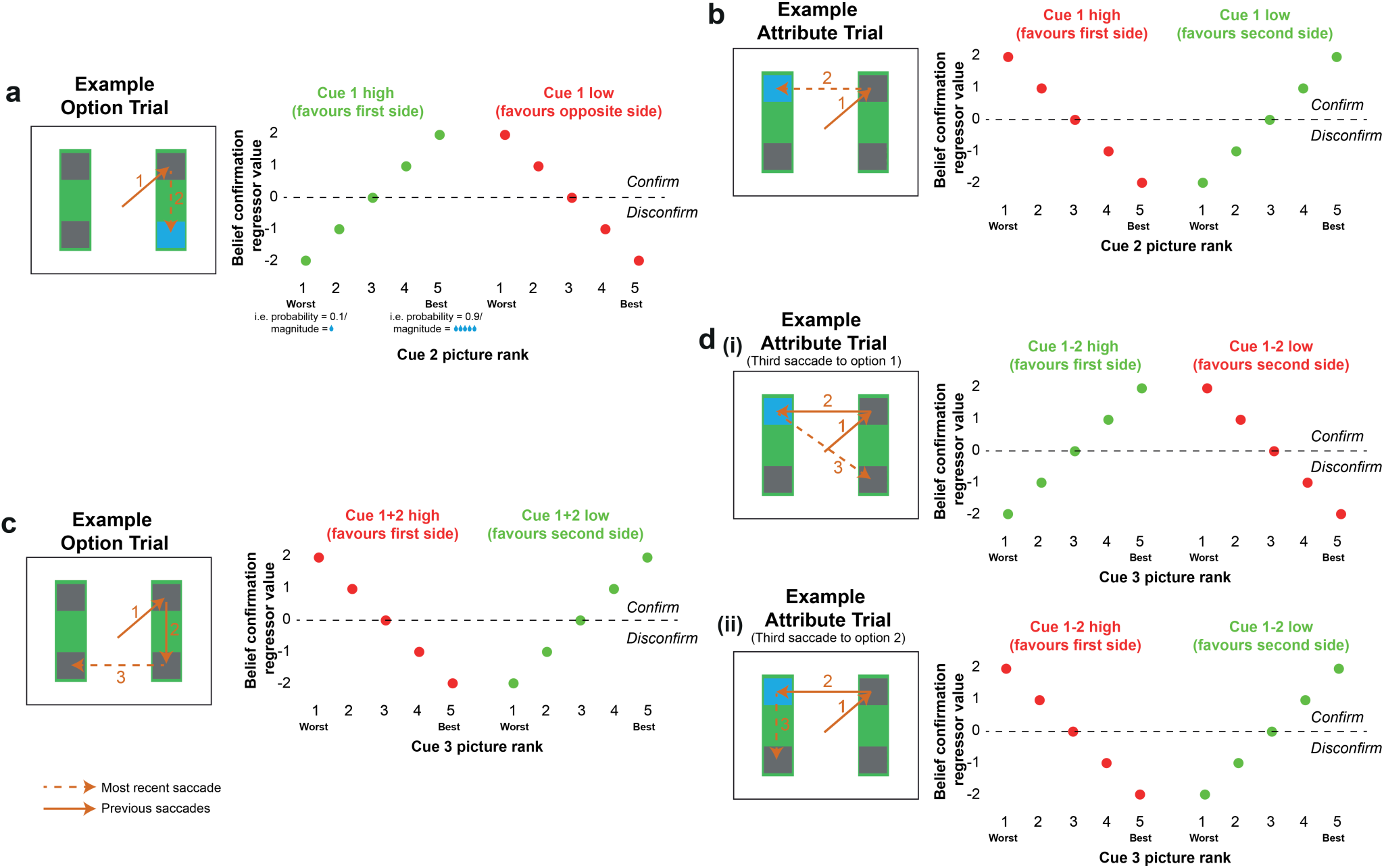
Design of ‘Belief confirmation regressors’ (i.e. EV13 – EV16 in General Linear Model). Each of the four EVs is depicted by a different panel, and refers to a different trial type/timepoint through the trial. Crucially, however, the interpretation of the four EVs is very similar. Whenever the evidence presented to the subject thus far suggests that the *currently attended* side should be chosen (green dots), then ‘belief confirmation’ scales *positively* with value. Whenever the evidence suggests that the *unattended* side should be chosen (red dots), then ‘belief confirmation’ scales *negatively* with value. Note that all four regressors were thus orthogonal to *currently attended* value (see **Extended Data Figure 10**). (a) EV13, reflecting belief confirmation at second saccade of option trials. (b) EV14, reflecting belief confirmation at second saccade of attribute trials. (c) EV15, reflecting belief confirmation at third saccade of option trials. (d) EV16, reflecting belief confirmation at third saccade of attribute trials (depending upon whether subjects’ third saccade was to (i) side 1, or (ii) side 2).

**Extended Data Figure 9.**
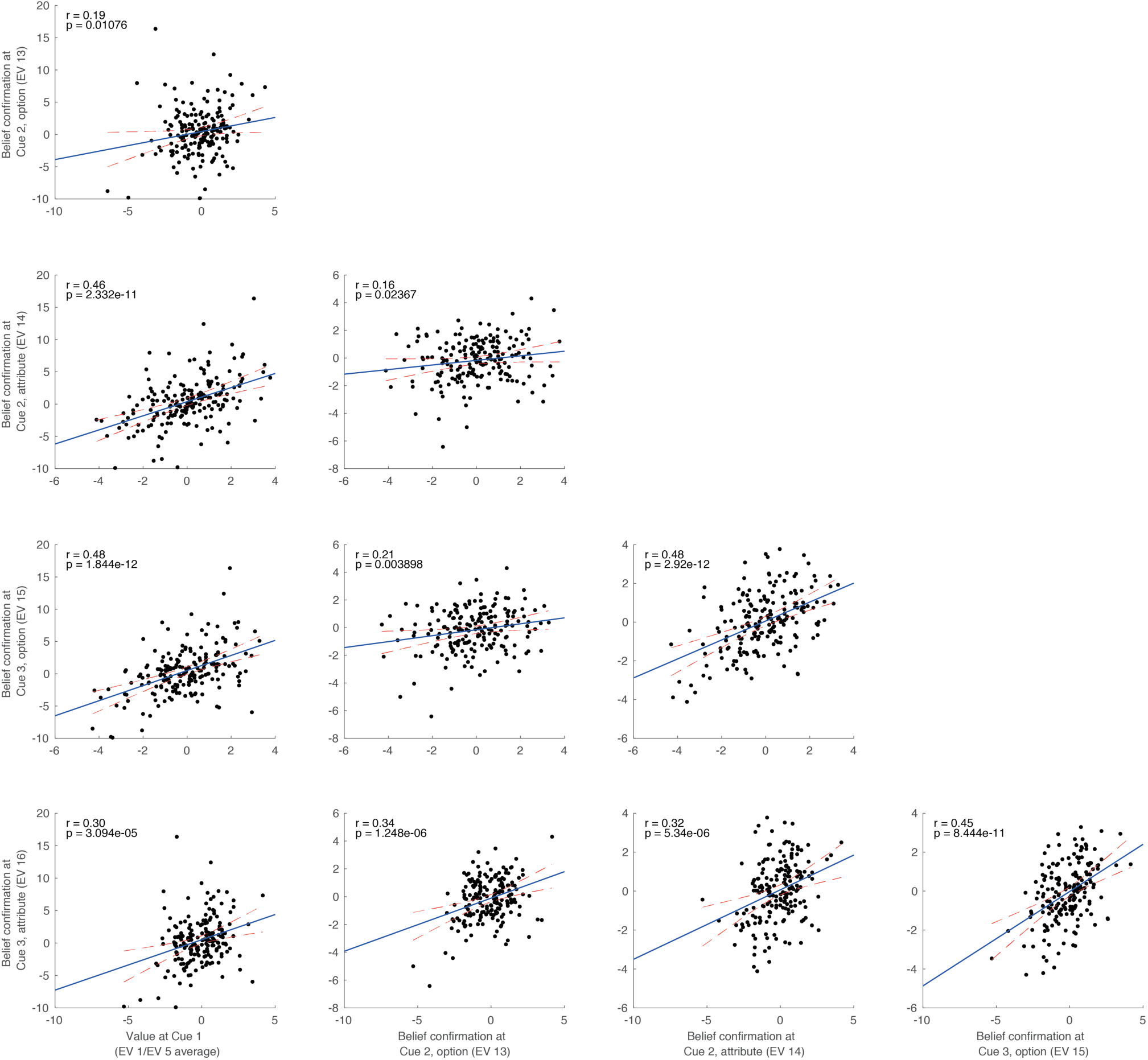
ACC has a robust belief confirmation signal across different cues and trial types. Parameter estimates for all four ‘belief confirmation’ regressors (at Cues 2/3 on both option and attribute trials) are positively correlated with each other across the ACC neural population. They are also positively correlated with value coding at cue 1. The correlation coefficients from each of these plots are also shown in **main Figure 4a**, middle plot.

**Extended Data Figure 10.**
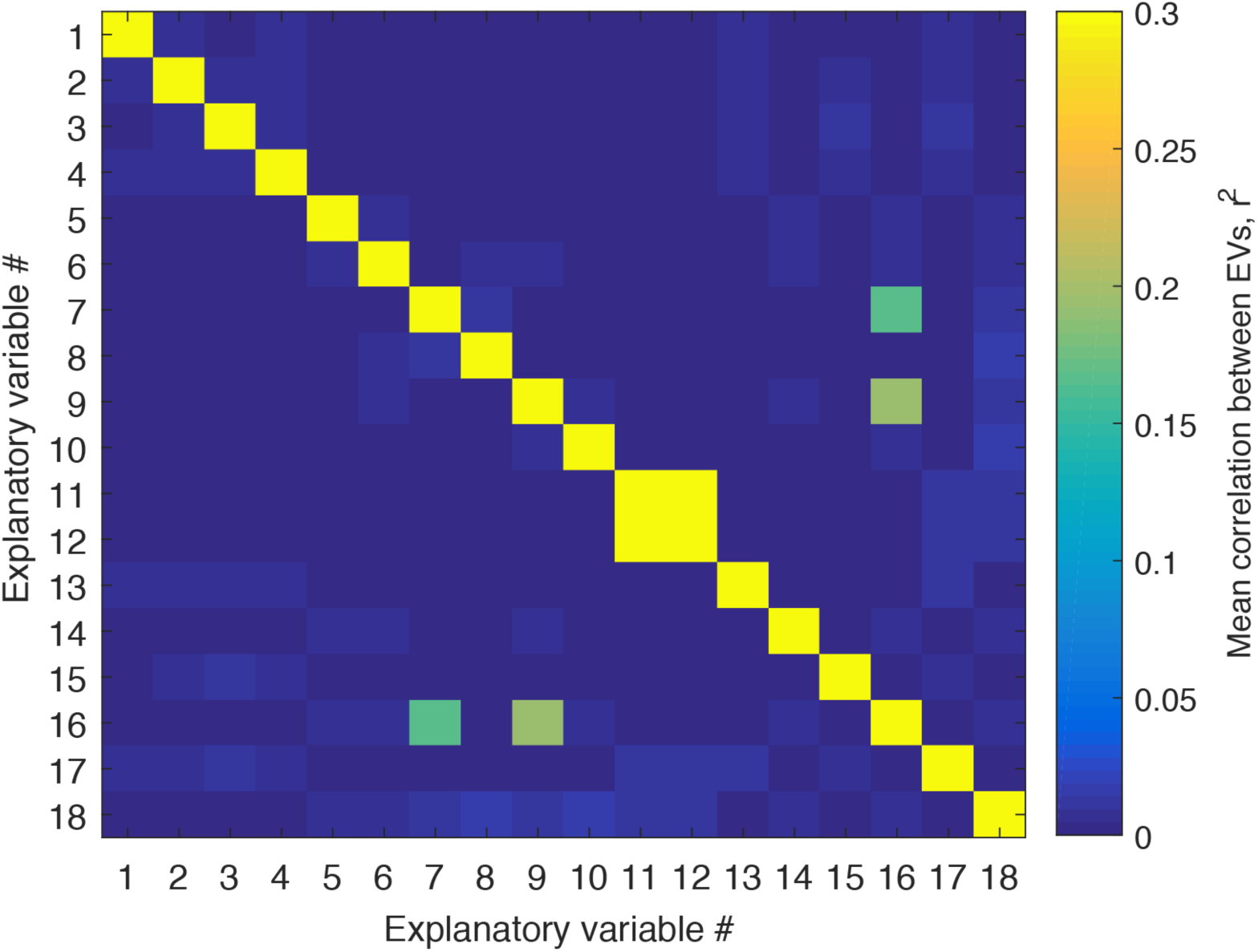
Mean correlation between EVs in General Linear Model. Note that most EVs of interest (1-6, 13-18) are decorrelated from one another by design, with the exception of EV16 (whose value depends upon where the subject looked at Cue 3, and this saccade depends systematically upon the relative value of cue 1 and cue 2 (see **supplementary note 1** for discussion)). EVs 10/11 are indicator variables for trial type.

**Extended Data Movie 1.** This movie depicts the temporal evolution of representational similarity around the time of Cue 1 fixation (see main **Figure 2)**. Each movie frame represents average firing rates of +/- 100ms around the timebin of interest. The bottom panels show the evolution of the template-based regression presented in main **Figure 2**.

**Extended Data Movie 2**. This movie shows the relationship between ‘belief confirmation’ and ‘left/right action selection’ population subspaces, both of which show ramping prior to action selection in anterior cingulate cortex.

## Methods

### Subjects

Two adult male rhesus monkeys (*Macaca mulatta*), M and F, were used as subjects and weighed 7-10kg at the time of neuronal data collection. We regulated their daily fluid intake to maintain motivation on the task. All experimental procedures were approved by the Local Ethical Procedures Committee and carried out in accordance with the UK Animals (Scientific Procedures) Act.

### Behavioural Protocol

Subjects sat head restrained in a behavioural chair facing a 19” computer monitor placed approximately 57cm away from the subjects’ eyes. The height of the screen was adjusted so that the centre of the screen aligned with neutral eye level for the subject. A voltage gating joystick (APEM Components, UK) was placed in front of the subject out of his line of sight and was used to make manual responses during the task. Eye position and pupil tracking was achieved using an infrared camera (ISCAN ETL-200) sampled at 240Hz.

The behavioural paradigm was run using the MATLAB based toolbox MonkeyLogic (http://www.monkeylogic.net/, Brown University, USA)^31-33^. All joystick and eye position was relayed to MonkeyLogic and for use online during the task and also recorded by MonkeyLogic at 1000Hz. Juice delivery was achieved by using a precision peristaltic (ISMATEC IPC) to pump juice to a spout placed at the lips of the subject. Subject M was given dilute (50%) apple juice while Subject F drank dilute (50%) mango juice.

Subjects were taught the value of a set of 10 isoluminant pictures cues pertaining to either magnitude or probability value (see *Task* for further details) using secondary conditioning on a separate day preceding data acquisition. This set of cues was then used for the following 1-4 recording sessions at which point a new set of cues would be taught to the subject. In total Subject M learnt 13 separate sets of cues, while Subject F learnt 11 sets.

### Task

A representation of the task structure is shown in main **Figure 1a**. Subjects initiated the trial by maintaining saccadic fixation on the centre of the screen and central fixation of the joystick for 500ms. Once this was achieved two options were presented on the screen (left and right of centre). Each option consisted of two pre-learned picture cues assigned to two different value attributes, probability of reward (10%, 30%, 50%, 70%, 90%) and magnitude of juice reward (0.15AU, 0.35AU, 0.55AU, 0.75AU, 0.95AU). The cues were uniformly sampled (with replacement, i.e. it was sometimes the case that the same cue would appear on both options). Reward magnitude (volume) was varied by manipulating the length of time a reward pump was driven, and the absolute values (i.e. reward time) associated with each stimulus varied slightly between subjects. Importantly, all four picture cues were covered up by grey squares with the exception of one which was covered by a blue square. The blue square informed the subject the location of a required saccade. Once the subject made a saccade and fixated the blue square, the blue square was replaced by the picture cue, which the subject was required to continuously fixate for 300ms. If continuous fixation was not achieved within 1200ms the trial was aborted and the subject received a short timeout. Once this fixation period was achieved, the cue was covered and a second blue square was presented, indicating the location of the required second saccade. The position of this blue square indicated to the subject the type of trial being experienced. If the blue square was for the second cue of the same option subjects were in an ‘Option trial’, whereas if the blue square was for the same attribute cue of the second option then this was an ‘Attribute trial’. Selection of trial types was pseudorandom. Once the subject made a saccade to the blue square, the blue square was replaced by the picture cue, and the subject was again required to maintain fixation of the second cue for 300ms. After this point, the subjects were relatively unconstrained. The two remaining unexplored locations were now replaced by blue squares. The subject could either choose an option using a joystick movement (left/right) based on the value of the currently known information, or saccade to one or both of the remaining cues (in any order) as they wanted (with the 3^rd^ cue requiring 300ms of fixation before subjects could saccade and uncover the information of the 4^th^ cue) before making a choice. Importantly, however, they were prevented from viewing any cue that they had already seen. Once a response was made all four cues were uncovered (for 500ms for Subject F and 1000ms for Subject M), after which juice reward feedback was given with the probability and reward magnitude chosen by the subject.

Note that the position of the probability/magnitude cues were counterbalanced across trials – i.e. on half of all ‘Option’/’Attribute’ trials, the probability cues would appear on the top row, and on the other half of trials the magnitude cues would appear on the top row. Attribute locations also corresponded across options (i.e. if probability was on the top row for the left option, it would also be on the top row for the right option). Additionally, the location of the first cue was counterbalanced across all four possible spatial locations across trials.

‘Option’ and ‘Attribute’ trials were pseudorandomly interleaved during blocks of 50 trials. Between each of these blocks subjects were presented with a block of 25 trials, where all four picture cues were presented immediately (so called ‘Simultaneous’ trials). Data from these trials will be discussed in a separate publication.

### Neuronal Recordings

Subjects were implanted with a titanium headpositioner for restraint, and then subsequently implanted with two recording chambers which were located using pre-operative 3T MRI and stereotactic measurements. Post-operatively we used gadolinium attenuated MRI imaging and electrophysiological mapping of gyri and sulci to confirm chamber placement. The centre of each chamber along the anterior-posterior (AP) coordinate plane was as follows; Subject M: left: AP 30.5, right: AP 33, Subject F: left: AP 34, right: AP 32.5. The chambers were angled along the medial-lateral plane to target different frontal regions (see **Extended Data Figure 1**). Craniotomies were then performed inside each chamber to allow neuronal recordings.

During each recording session, neuronal activity was measured using tungsten microelectrodes (FHC Instruments, Bowdoin, USA) that were lowered into the brain through a grid using using custom-built manual microdrives or chamber-mounted motorized microdrives (FlexMT; AlphaOmega Inc.). During a typical recording session 8-24 electrodes were lowered bilaterally into multiple target regions until well-isolated neurons were found. Neuronal data was recorded at 40kHz using a Plexon Omniplex system (Dallas, USA). Single unit isolation was achieved with manual spike sorting, using Plexon Offline Sorter (Dallas, USA). We randomly sampled neurons; no attempt was made to select neurons based on responsiveness. This procedure ensured an unbiased estimate of neuronal activity thereby allowing a fair comparison of neuronal properties between the different brain regions.

We recorded neuronal data from three target regions: anterior cingulate cortex (ACC), dorsolateral prefrontal cortex (DLPFC), and orbitofrontal cortex (OFC). We considered ACC to be the entire dorsal bank of the anterior cingulate sulcus from AP 27-37. Our LPFC recordings spanned both dorsal and ventral banks of the principal sulcus but were concentrated towards the former. All neurons recorded lateral to the medial orbital sulcus and medial to the lateral orbital sulcus was considered OFC. Some recordings were also made in ventromedial prefrontal cortex (VMPFC), but these will be discussed in a separate publication. We used the gadolinium-enhanced MRI along with electrophysiological observations during the process of lowering each electrode to estimate the location of each recorded neuron and produce a histological map of the neuronal population (see **Extended Data Figure 1**).

### Representational similarity analysis (RSA) at Cue 1 presentation (Figure 2 and Extended Data Figures 3/5, Extended Data Movie 1)

To calculate the representational similarity matrices shown in **Figure 2a-c**, we first calculated the average firing rate for each neuron for each of the 20 conditions of interest: when the lowest to highest probability cue was presented on the left at Cue 1, when the lowest to highest magnitude cue was presented on the left at Cue 1, lowest to highest probability cue on right, and lowest to highest magnitude cue on right. This firing rate was computed between 100ms and 500ms after Cue 1 onset. We then normalised across these 20 conditions, subtracting the mean and dividing by the standard deviation.

Repeating this for every neuron yielded a matrix with dimensions (neurons^∗^20). For two conditions (*i*, *j*), we computed the correlation coefficient across neurons between row *i* and row *j* of this matrix, which is plotted in element (*i*,*j*) of the representational similarity matrix. For **Extended Data Movie 1**, we repeated the same procedure on sliding windows of +/- 100ms from the timepoint of interest.

### RSA template-based regression (Figure 2d-h)

We used multiple linear regression to assess the contribution of several potential ‘template’ neural codes to the RSA matrices within each region. Each of the 400 elements of each region’s RSA template was explained using the following regression model:

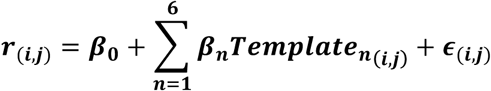

Where ***r*** denotes the correlation coefficient matrix computed using RSA, and there are six ‘template’ matrices onto which the RSA matrix is regressed. We estimated ***β*_*0*-*6*_** using ordinary least squares, minimising the sum of squared residuals ***ε***. The six template matrices were as follows:

Template 1: Identity matrix – accounting for all RSA matrices being 1 when element *i*=element *j*

Template 2 **(Figure 2d)**: ‘Spatial attention’ – accounting for representational similarity between cues presented on the same side, but dissimilarity between cues on opposite sides (1 if *i*<=10 and *j*<=10, 1 if *i*>=11 and *j*>=11, 0 elsewhere)

Template 3 **(Figure 2e)**: ‘Stimulus identity’ – accounting for representational similarity between the same stimulus being presented on left/right options (1 where |*i*-*j*| = 10, 0 elsewhere)

Template 4 **(Figure 2f)**: ‘Attended value’ – accounting for representational similarity between similarly valued items and representational dissimilarity between dissimilarly valued items (*ranked value (i)*^∗^*ranked value (j)*, where ranked value is -2 for the lowest ranked stimulus within an attribute (i.e. 10% probability, 15% maximal reward magnitude), -1 for the 2^nd^ lowest ranked (30% probability, 35% maximal reward magnitude), 0 for the median ranked (50 % probability, 55% maximal reward magnitude), 1 for the 2^nd^ highest ranked (70% probability, 75% maximal reward magnitude), 2 for the highest ranked (90% probability, 95% maximal reward magnitude))

Template 5 **(Figure 2g)**: ‘Left/right value’ – interaction of template 2 and template 4

Template 6 **(Figure 2h):** ‘Accept/reject’ - accounting for representational similarity between cues that might lead to ultimately accepting the current alternative (good items similar to other good items; bad items similar to other bad items), and representational dissimilarity between dissimilar items in terms of acceptance/rejection (*sign* of *attended value* template)

For the middle panels in **Figure 2d-h** this model was estimated on RSA matrices from 100-500ms post-stimulus, as in **Figure 2a-c**; for the bottom panels of **Figure 2d-h** it was performed on sliding windows of +/- 100ms from the timepoint of interest, as in **Extended Data Movie 1**. In these panels we plot the coefficient of partial determination (CPD) for each regressor across time, which is defined for EV X_i_ as follows:

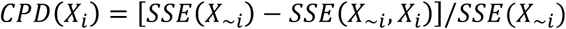

where *SSE(X)* refers to the sum of squared errors in a GLM that includes a set of EVs *X*, and *X_~i_* is a set of all the EVs included in the full model except *X_i_*^14,34^.

### Statistical inference on RSA template-based regression model

We tested the significance of each template within each region by computing the T-statistic for each *β* coefficient (i.e. *β̂_n_*/*σ̂_n_*, where *σ̂_n_* denotes the standard errors of each coefficient estimate). We compared differences between regions by computing F-statistics equivalent to a one-way ANOVA (see https://fsl.fmrib.ox.ac.uk/fsl/fslwiki/FEAT/UserGuide#ANOVA:_1-factor_4-levels for example). Importantly, however, when calculating these statistics on a correlation matrix, they may not be parametrically distributed in the null distribution (due to observations not being independently and identically distributed). To overcome this, we built a non-parametric null distribution for each test of interest, by permuting the identities of the 20 cues (i.e. values 1-5 on probability/magnitude, left/right), recomputing the RSA matrix, and rerunning the regression. We then computed the T-statistics and F-statistics on this permuted data, and compared the true statistics to the permuted null distribution to obtain *p*-values^35^. We performed 10,000 permutations.

### General linear model (GLM), underlying analyses in Figures 3 and 4

For the population analyses shown in **Figures 3** and **4**, we first estimated a general linear model on the firing rate of each individual neuron, timelocked with respect to Cue 1 presentation, Cue 2 presentation, Cue 3 presentation, and joystick movement (response). Each neuron’s firing rate was explained using a GLM containing 18 explanatory variables (EVs), detailed below, estimated using ordinary least squares. Note that *EVs 1*-*6* are critical for the analyses shown in **Figure 3** and **Extended Data Figures 6-7**, *EVs 13*-*16* are critical for the analyses shown in **Figure 4a-c** and **Extended Data Figure 9**, and *EVs 17*-*18* are critical for the analyses shown in **Figure 4d**.

EV 1 captured the linear effect of changing the first attended cue’s value from the lowest value to highest value, collapsing across probability and magnitude cues, selectively on ‘option trials’. Specifically, if the lowest ranked probability/magnitude item was presented they were valued -2; if the second lowest ranked item was presented -1; third lowest, 0; second highest, 1; highest, 2.

EVs 2-4 were similar to EV 1, but reflected the second, third and fourth attended cue’s value respectively (for option trials only). On trials where the third or fourth cue was not attended on an option trial (because the subject responded without sampling all cues), the corresponding EVs were valued 0.

EVs 5-6 were similar to EVs 1-2, but reflected the first and second attended cue’s value respectively for ‘attribute trials’ only. EVs 7-8 were similar to EVs 3-4, but reflected the third and fourth attended cue’s value on attribute trials where the subject saccaded diagonally back to the first side of the screen (0 on vertical saccade trials), whereas EVs 9-10 reflected the third and fourth attended cue’s value on attribute trials where the subject saccaded vertically to the second side of the screen (i.e. 0 on diagonal saccade trials). Note that there is no need to split option trials by third saccade direction, as unlike in option trials the third saccade is always to the second side of the screen.

EV 11 was an indicator variable for option trials (1 on option trials, 0 otherwise); EV 12 was an indicator variable for attribute trials (1 on attribute trials, 0 otherwise). Note that EVs 11 and 12 sum to produce a constant term, thereby capturing variation in the mean firing rate of the cell across time.

EVs 13-16 were variables that all captured the extent to which the Cue value observed at Cue 2 and Cue 3 were *consistent* (belief confirmation) or *inconsistent* (belief disconfirmation) with the currently held belief as to which option would be rewarded. They are described below, but for clarity, they are also depicted in **Extended Data Figure 8**. Two key points are pertinent: (a) by design, all four EVs were largely *orthogonal to the value of Cue 1*, *Cue 2 and Cue 3* (although see note on EV 16 below); (b) they each rely upon different cues and different trials, and so are *orthogonal to each other* by design.

EV 13 **(Extended Data Figure 8a)** was the same as EV 2 – i.e. the value of cue 2 on option trials – but crucially, it was multiplied by 1 whenever the value of the first cue was *greater* than the average value (i.e. best or second best picture cues), multiplied by - 1 whenever the value of the first cue was *lower* than the average value (i.e. worst or second worst picture cue), and multipled by 0 whenever it was of average value (middle picture cue). EV 13 therefore was positively signed whenever Cue 2 was consistent with Cue 1 (e.g. low-valued cue followed by another low-valued cue, or high-valued cue followed by another high-valued cue).

EV 14 **(Extended Data Figure 8b)** was the same as EV 6 – i.e. the value of cue 2 on attribute trials – but was multiplied by 1 when the first cue’s value was *lower* than average, by -1 whenever the first cue’s value was *higher* than average, and by 0 when cue 1 was of average value. Again, this meant that EV 14 was positively signed whenever it was consistent with Cue 1 (e.g. low-valued cue on the left followed by high-valued cue right both favour the right action, or high-valued cue on the left followed by low-valued cue on the right both favour a left action).

EV 15 **(Extended Data Figure 8c)** was the same as EV 3 – i.e. the value of cue 3 on option trials – but was multiplied by 1 whenever the first and second cue were lower than average value, by -1 whenever the first and second cue were higher than average value, and by 0 when the first and second cue were of average value.

EV 16 **(Extended Data Figure 8d)** was similarly defined to EVs 7 and 9 – i.e. the value of cue 3 on attribute trials – but crucially relies upon an interaction of the relative value of the first and second cue, and which side the subject decided to attend with the third saccade. On trials where the subject’s third saccade was diagonal back to option 1, it was EV 7 multiplied by 1 when (Cue 1 value>Cue 2 value), multiplied by -1 when (Cue 2 Value>Cue 1 value), and multiplied by 0 when (Cue 1 value=Cue 2 value). On trials where the subject’s third saccade was vertical within option 2, it was EV 9 multiplied by 1 when (Cue 2 value>Cue 1 value), multiplied by -1 when (Cue 1 Value>Cue 2 value), and multiplied by 0 when (Cue 1 value=Cue 2 value). Note that because subjects’ decision whether to make a third saccade to the same side as option 1 relied upon the relative value of Cue 1 and Cue 2 (see **supplementary note 1**), there existed some positive correlation between EV16 and EVs 7 and 9 (mean r^2^ of 0.167 and 0.194 respectively, see **Extended Data Figure 10**). Nevertheless, including all three EVs together in the GLM directly controls for this correlation with value, by partialling out any variance that can be attributed to EVs 7 or 9 from the parameter estimate for EV 16.

EV 17 was defined in terms of action selectivity on option trials. It was valued 1 on option trials where the subject chose left, -1 on option trials where the subject chose right, and 0 on attribute trials.

EV 18 was defined in terms of action selectivity on attribute trials. It was valued 1 on attribute trials where the subject chose left, -1 on attribute trials where the subject chose right.

We estimated this multiple regression model on neuronal firing rate in sliding 200ms bins, stepped in 10ms time-windows, from 100ms pre-cue to 500ms post-cue (when stimulus-locked), or from 500ms pre-response to 100ms post-response (when response-locked). We excluded trials where subjects viewed fewer than 3 cues from this analysis.

### Peri-stimulus correlation and cross-correlation of parameter estimates from GLM2 (Figure 3/Extended Data Figures 6-7)

Once the model in the previous section was estimated for each neuron, we then correlated, *across neurons*, T-statistics associated with parameter estimates for different EVs. This allowed us to examine how population subspaces encoding different variables related to each other, at various timepoints through the trial. Note that in one case (**main fig 3b**) we collapse across parameter analyses from option and attribute trials for clarity. Parameter estiamtes in **Figure 3b-f** were taken from 300ms post-stimulus, whereas in **Figure 3g-j** they were repeated on all possible combinations of time-points to produce cross-correlation matrices of parameter estimates.

### Projection of ACC activity onto belief confirmation/chosen response subspaces (Figure 4 and Extended Data Figure 9, Extended Data Movie 2)

To identify whether there was a stable subspace representing ‘belief confirmation’ in each brain region (**Extended Data Figure 9**), we investigated whether the parameter estimates for all four regressors that captured belief confirmation in our GLM were correlated. The parameter estimates used were EV 13, 300ms after Cue 2 onset; EV 14, 300ms after Cue 2 onset; EV 15, 300ms after Cue 3 onset; EV 16, 300ms after Cue 3 onset. We also asked whether this subspace was similar to the subspace for Cue 1 value (i.e. EV1 + EV5, 300ms after Cue 1 onset), based on the idea that Cue 1 ‘value’ responses in ACC are better conceived in terms of belief confirmation about accepting or rejecting the first attended cue (cf. results in **Figure 2c, 2h)**.

Once this stable subspace was identified in ACC (see **Extended Data Figure 9**), we asked how activity in this subspace evolved in trials where the subject took different lengths of time to make his final choice response (**Figure 4** and **Extended Data Movie 2**). For each neuron, we split trials into five separate bins depending upon response time from Cue 1 onset, and averaged neuronal firing for these different trial types. For each bin, this yielded a matrix with dimensions *time*^∗^*neurons*.

To examine activity within different subspaces, we then regressed this matrix onto a projection matrix composed of two key ‘weights’ per neuron, i.e. T-statistics of contrasts of parameter estimates of interest, estimated from the GLM. This projection matrix therefore had dimensions *neurons*^∗^*(2 PEs).* The two contrasts of interest were:

1. The average parameter estimates for belief confirmation, i.e. EV 13, 300ms after Cue 2 onset; EV 14, 300ms after Cue 2 onset; EV 15, 300ms after Cue 3 onset; EV 16, 300ms after Cue 3 onset;
2. The average parameter estimates for left vs. right action selection, i.e. EV 17 and EV 18, 200ms prior to response onset;

Regressing the time^∗^neurons matrix onto the neurons^∗^(3 PEs) gives rise to the sliding analysis that is shown in **Figure 4**. In **Figure 4b/c**, we plot the stimulus-locked and response-locked parameter estimates for contrast 1 respectively, reflecting the population activity in the belief confirmation subspace for trials of different length. In **Figure 4d**, we plot the response-locked parameter estimates for contrast 2, reflecting population activity in the left/right action selection subspace in trials of different length. In both cases, we baseline corrected subspace activity to the time of Cue 1 onset +/- 50ms. **Extended Data Movie 2** provides a representation of how activity in both of these subspaces progresses during the course of the trial.

Crucially, we avoided using the same data for estimating different neurons’ weights in the projection matrix as for plotting population activity. To achieve this, we first split the data into odd and even trials; we estimated the projection matrix weights using the GLM on the odd trials, and projected these weights onto firing rates on the even trials; we then repeated the same process with even trials for GLM estimation and odd trials for projection; finally, we averaged subspace activity together across odd and even-trial analyses.

### Data availability statement

Data and analysis scripts to reproduce figures from the paper will be made publicly available for download on the CRCNS repository (http://crcns.org/) upon publication.

## Acknowledgements

L.T.H. was supported by a Henry Wellcome Fellowship from the Wellcome Trust (098830/Z/12/Z), and a NARSAD Young Investigator Grant from the Brain and Behavior Research Foundation. N.M was supported by funding from the Astor Foundation, Rosetrees Trust and Middlesex Hospital Medical School General Charitable Trust. A.D.B. was supported by a PhD studentship from the MRC. B.M. was supported by the Fundação para a Ciência e Tecnologia (scholarship SFRH/BD/51711/2011). T.E.J.B. was supported by a Wellcome Trust Senior Research Fellowship (WT104765MA) and funding from the James S McDonnell Foundation (JSMF220020372). S.W.K. was supported by a Wellcome Trust New Investigator Award (096689/Z/11/Z).

